# ReQTL – an allele-level measure of variation-expression genomic relationships

**DOI:** 10.1101/464206

**Authors:** Liam Spurr, Nawaf Alomran, Piotr Słowiński, Muzi Li, Pavlos Bousounis, Qianqian Zhang, Justin Sein, Keith A. Crandall, Krasimira Tsaneva-Atanasova, Anelia Horvath

## Abstract

**Motivation:** By testing for association of DNA genotypes with gene expression levels, expression quantitative trait locus (eQTL) analyses have been instrumental in understanding how thousands of single nucleotide variants (SNVs) may affect gene expression. As compared to DNA genotypes, RNA genetic variation represents a phenotypic trait that reflects the actual allele content of the studied system. RNA genetic variation can be measured at expressed genome regions, and differs from the DNA genotype in sites subjected to regulatory forces. Therefore, assessment of correlation between RNA genetic variation and gene expression can reveal regulatory genomic relationships in addition to eQTLs.

**Results:** We introduce ReQTL, an eQTL modification which substitutes the DNA allele count for the variant allele frequency (VAF) at expressed SNV loci in the transcriptome. We exemplify the method on sets of RNA-sequencing data from human tissues obtained though the Genotype-Tissue Expression Project (GTEx) and demonstrate that ReQTL analyses show consistently high performance and sufficient power to identify both previously known and novel molecular associations. The majority of the SNVs implicated in significant cis-ReQTLs identified by our analysis were previously reported as significant cis-eQTL loci. Notably, trans ReQTL loci in our data were substantially enriched in RNA-editing sites. In summary, ReQTL analyses are computationally feasible and do not require matched DNA data, hence they have a high potential to facilitate the discovery of novel molecular interactions through exploration of the increasingly accessible RNA-sequencing datasets.

**Availability and implementation:** Sample scripts used in our ReQTL analyses are available with the Supplementary Material (ReQTL_sample_code).

**Supplementary Information:** Re_QTL_Supplementary_Data.zip

## 1. Introduction

Quantitative trait loci (QTL)-based approaches have served as a major tool to uncover genetic variants regulating phenotypic features. In recent years, QTL approaches have been successfully applied to a variety of molecular traits, including gene expression (eQTL), splicing (sQTL), protein expression (pQTL), methylation (meQTL), chromatin accessibility (chQTL/caQTL) and histone modification (hQTL/cQTL) (Albert and Kruglyak, 2015; Atak *et al.*, 2013; Aguet *et al.*, 2017a; Weiser *et al.*, 2014; Li *et al.*, 2015; Brandt and Lappalainen, 2017; Odhams *et al.*, 2017; Ko *et al.*, 2017; Winter *et al.*, 2018; Heinig, 2018; De Almeida *et al.*, 2018). To correlate genetic variants with the trait of choice, all these methods utilize the genotypes obtained through DNA analysis for each SNV locus.

Compared to DNA, RNA molecules carry and present genetic variation in a related yet distinct manner, where the differences reflect regulatory or selective forces. The two most common mechanisms causing differences between DNA and RNA alleles at SNV loci are allele specific expression (ASE) and RNA editing. ASE results in asymmetric RNA content derived from two chromosomes, and is frequently caused by cis-acting genetic variants affecting the function, structure, stability, or the speed of transcription of the RNA molecule (Chess, 2016). RNA editing is a post-transcriptional modification, which, similarly to the allele-specific expression, can result in different RNA function, stability, or sequence, including protein altering changes and motifs recognizable by RNA-binding molecules(Eisenberg and Erez Y. Levanon, 2018; Guo *et al.*, 2018). Therefore, assessment of correlation between RNA genetic variation and gene expression can reveal molecular relationships in addition to those identifiable through eQTLs using genotypes. Moreover, RNA genetic variation represents a phenotypic trait that reflects the actual allele content of the studied system.

For diploid genomes, a commonly used measure for the RNA genetic variation is the expressed variant allele frequency, VAF_RNA_, which can be estimated from the RNA-sequencing data (VAF_RNA_ = **n**_var_ / (**n**_var_ + **n**_ref_)), where **n**_var_ and **n**_ref_, are the variant and reference read counts (29267927). In contrast to the categorical DNA-allele count of 0, 1 and 2, VAF_RNA_ is a continuous measure, which allows for precise allele quantitation. Furthermore, as compared to DNA genotypes, VAF_RNA_ reflects allele-level expression regulation, including imprinting, expression regulation mediated by RNA-binding molecules, and RNA-editing.

Herein, we propose a method to assess variation-expression relationships based on VAF_RNA_-derived information on genetic variation; we term the method ReQTL (**R**NA-**eQTL**). We have based our model on the underlying assumption of eQTLs: if a given variant affects the expression of a given gene, the expression of this gene scales with the number of alleles harboring the variant of interest. This assumption intuitively implies both DNA-mediated effects (i.e., effects mediated via DNA-binding molecules), and effects resulting from solely RNA-mediated interactions. For the majority of the expressed SNV positions, the RNA-transcription is expected to scale with the DNA-allele count. Hence, ReQTL analysis captures both DNA-defined variation-expression correlations, and RNA-exclusive variation-expression relationships such as those involving RNA editing. ReQTL analyses can be run directly on eQTL-built computational platforms. The proposed pipeline (Figure 1) employs publicly available packages for processing of sequencing data, and sample code for our ReQTL-specific data transformation is available in the Supplementary Files (ReQTL_sample_code). We exemplify ReQTL analysis using the popular software Matrix eQTL (Shabalin, 2012) on RNA-sequencing data obtained from the Genotype-Tissue Expression (GTEx) project (www.gtexportal.org, phs000424.v7), from three different tissue types: Nerve-Tibial, Skin-Sun-Exposed (lower leg), and Skin-Not-Sun-Exposed (suprapubic). We identify both common and tissue-specific ReQTLs, as well as both known and novel molecular relationships. In our data, trans-ReQTLs were significantly enriched in RNA-editing sites.

**Figure 1.**
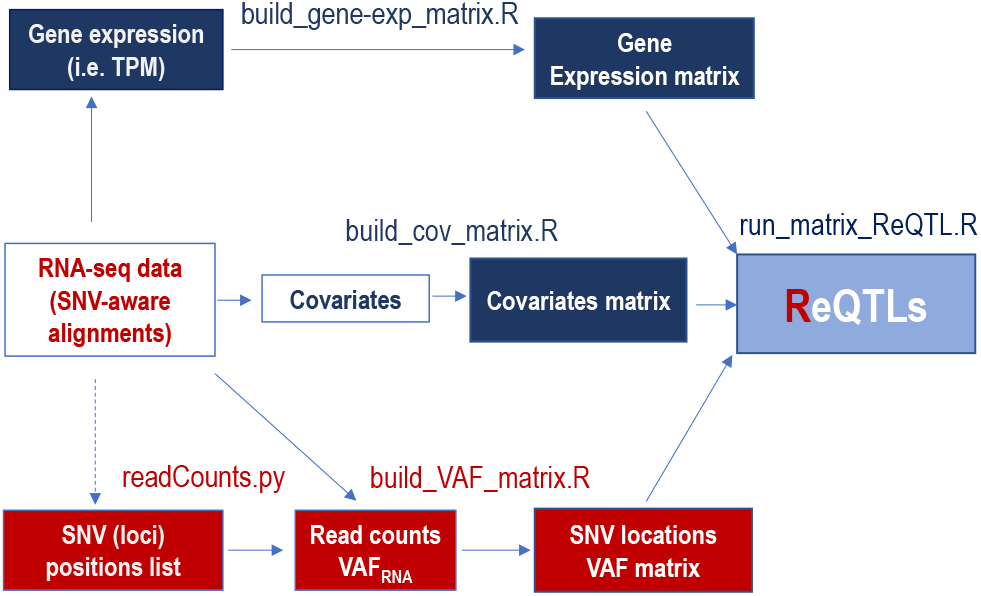
Major steps of the ReQTL analyses (differences from eQTL analysis are outlined in red). SNV-aware alignments are used to generate gene-expression data; TPM values are quantile transformed and used to generate gene-expression matrix (exemplified by build_gene-exp_matrix.R). Lists of genomic positions can be built using any custom set of positions of interest (i.e., dbSNP). Alternatively, lists of genomic positions can be generated through variant call and subsequent retainment of the unique variant genomic loci across the sample set. At each genomic position in the list, the reference and variant number of RNA-sequencing reads are counted from the alignments and used to estimate VAF_RNA_ in each individual sample from the set (readCounts.py from our lab’s read-Counts package). The VAF_RNA_ estimations are used to build VAF matrix (exemplified by build_VAF_matrix.R). Covariates can be accounted for by using approaches similar to the ones used in eQTL analyses (exemplified by build_cov_matrix.R). The three matrices are then used as input for Matrix eQTL (exemplified by run_matrix_ReQTL.R).

## 2. Methods

### 2.1. Samples

A total of 772 raw RNA-sequencing datasets from three different body sites – Nerve–Tibial (238 samples), Skin-Sun-Exposed, lower leg (286 samples), and Skin-Not-Sun-Exposed, suprapubic (248 samples) - were downloaded on 06/10/18 from the Database of Genotypes and Phenotypes (dbGaP, https://www.ncbi.nlm.nih.gov/gap). All the libraries were generated using non-strand specific, polyA-based Illumina TruSeq protocol, and sequenced to a median depth of 78 million 76-bp paired-end reads. The selection of tissue types was based on the availability of more than 200 samples, and consideration for assessment of both distinct (Nerve vs Skin) and related (Skin-Sun-Exposed vs Skin-Not-Sun-Exposed) tissue types.

### 2.2. Data processing

RNA-sequencing reads were aligned to the latest release of the human reference genome (hg38/GRCh38, Dec 2013) using HISAT2 (version 2.1.0) with an SNV-aware index covering over 12 million SNPs (Kim *et al.*, 2015). SNV-aware alignment is recommended to avoid mapping bias in the computation of the allele frequency. On the aligned RNA sequencing reads, we called variants using GATK (version 4.0.8.0) and retained positions in the genome reference index with high quality calls in the individual samples (QUAL>100, MQ>55 for further analysis (Van der Auwera *et al.*, 2013).

Within each tissue type, we combined the SNVs called across all samples into a list of unique SNV positions. We then estimated n_var_ and n_ref_, and computed VAF_RNA_ for each of the positions in the list in each of the individual samples. To do that, we used the module readCounts previously developed in our lab (http://github.com/HorvathLab/NGS/tree/master/readCounts, (Movassagh *et al.*, 2016). Briefly, readCounts employs the pysam Python module to assess the read counts at every SNV position of interest in each of the alignments (samples) from a studied group (i.e. tissue). ReadCounts then filters aligned reads based on alignment quality metrics including length, gaps and mapping quality, and tallies the remaining reads as having the expected reference or variant nucleotides. Importantly, for ReQTL analyses, we retained only positions covered by a minimum of 10 total sequencing reads in all of the samples from a studied group. Additionally, we excluded variants with a monoallelic or missing signal in more than 80% of the samples from a given tissue.

Gene expression was estimated from the alignments using Stringtie (version 1.3.4.) (Kim *et al.*, 2015), and TPM (transcripts per million) values were used for the ReQTL estimation. Pseudogenes were identified based on ensemble annotations (https://useast.ensembl.org/info/data/biomart/index.html), and ex-cluded from the analysis. Furthermore, within each tissue, we filtered out genes with TPM value < 1 in more than 80% of the samples. The TPM distribution was quantile-transformed using the average empirical distribution observed across all samples in the corresponding tissue (Aguet *et al.*, 2017a). For each gene, the TPM values were transformed to the quantiles of the standard normal distribution. The effects of unobserved confounding variables on gene expression were quantified using probabilistic estimation of expression residuals (PEER), with 25 PEER factors (Stegle *et al.*, 2012).

## 3. Results

### 3.1. ReQTL analysis

We performed the ReQTL analyses separately for the three tissues, using a linear regression model as implemented in the package Matrix eQTL (Shabalin, 2012). To account for covariates, we corrected for the top 25 PEER factors (Stegle *et al.*, 2012), reported age, race, sex, and the top ten VAF_RNA_ principal components (PCs). Lists of loci were generated based on the combined variation call in each tissue, after filtering for quality and frequency of the variant in the studied group. To be considered cis-ReQTL, a variant was required to reside within 1Mb (megabase) of the transcription start site of a gene. We retained for further analysis significant associations (FDR < 0.05) with a strong positive or negative correlation (Spearman’s rank correlation coefficient ρ ≥ 0.3 or ρ ≤-0.3).

### 3.2. Overall findings

Under the described settings, we identified 21,371 cis- and 2,280 trans-SNV loci implicated in ReQTLs using a significance level of FDR < 0.05 across the three tissues (Supplementary Tables 2 and 3, respectively). These SNVs participated in a total of 31,301 cis- and 17,800 trans-correlations (Supplementary Tables 4 and 5, respectively) with 4712 and 1842 unique cis- and trans-regulated genes (Supplementary Tables 6 and 7, respectively). Similar to the eQTL analyses, a high proportion of the effects were tissue-specific, with substantially higher overlap between the two related tissues (Skin-Sun-Exposed and Skin-Not-Sun-Exposed) and higher tissue specificity in the trans- (as compared to cis-) ReQTLs (Figure 2a). Quantile-quantile (QQ) plots are presented on Figure 2b.

**Figure 2.**
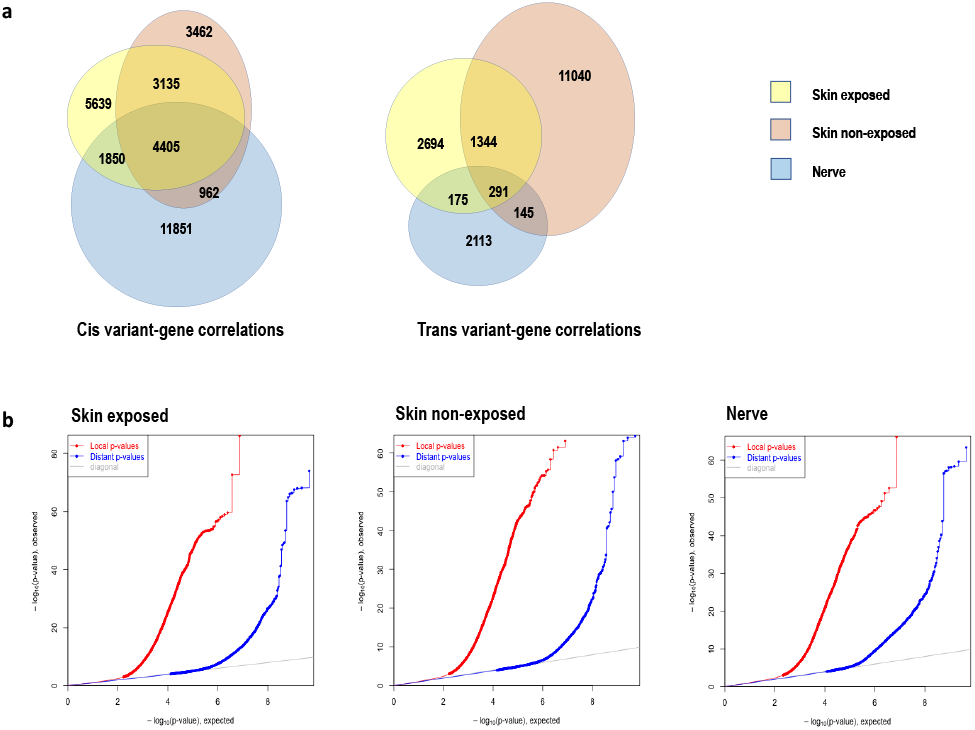
**a.** Tissue-specific and shared ReQTL correlations in cis (left) and trans (right) correlations between tissues. As expected, substantially higher overlap is seen in the cis-correlations. **b.** Quantile-quantile plot of −log10 p-values for local and distal ReQTLs in the three tissue types. To be considered a cis-correlation, an SNV and the transcription starting site of a gene needed to be within 10e6 bases. Similar to the eQTLs, higher numbers of significant correlations were found in cis-mode.

We observed three distinct correlation patterns (Figure 3). First, a substantial proportion of the correlation plots identified a pattern similar to an eQTL plot (Figure 3a). For these patterns, the VAF_RNA_ distribution appears to reflect to a large extent the genotype distribution. This type of correlation is likely to be explained by mechanisms similar to those underlying the eQTLs. The second type of pattern was related to the first, with VAF_RNA_ values of 0 or 1 corresponding to homozygous genotypes, and intermediate VAF_RNA_ values spread along the linear regression line (Figure 3b). These correlations are likely to reflect quantitative effects of variants on gene expression, where discrete increase/decrease in the frequency of particular alleles leads to gradual changes in the gene expression. The third type of ReQTLs showed a distinct distribution, where the individual measurements closely followed the association trend, with fewer VAF_RNA_ values of 0 or 1 (Figure 3c). These patterns were seen in the majority of the visually examined trans-ReQTLs (the top 1000 significant correlations of each tissue type). Further analysis of these ReQTLs showed that a substantial proportion of the involved loci overlap with known RNA-editing positions (see below).

**Figure 3.**
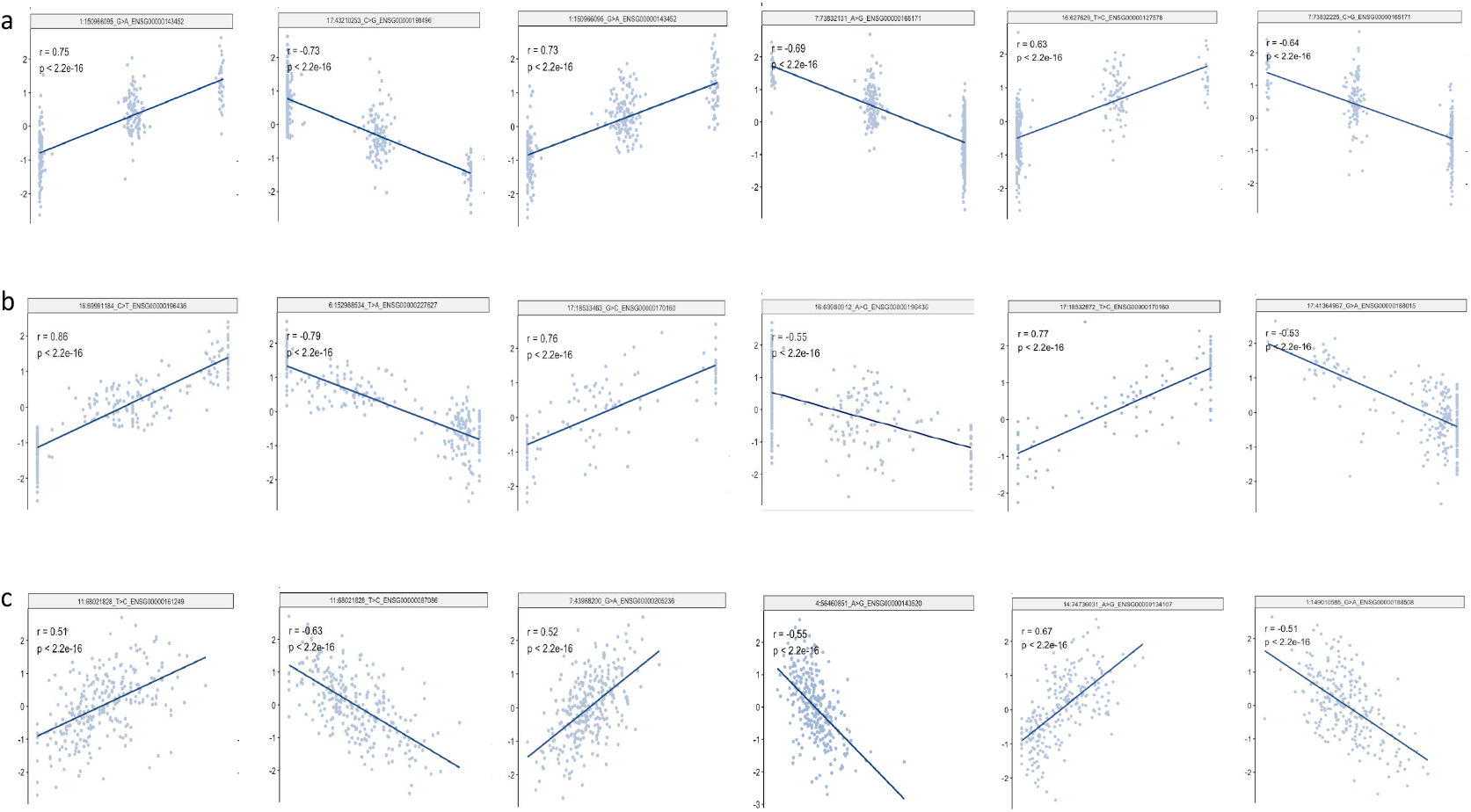
Correlation patterns identified by ReQTL analyses. **a.** eQTL-like patterns were seen in a large number of the correlations that overlap with significant eQTLs**. b.** A proportion of correlations show plots with non-extreme VAF_RNA_ values spread along the linear regression line, and VAF_RNA_ of 0 or 1 similar to the first pattern. **c.** A characteristic plot pattern with most of the VAF_RNA_ values spread along the linear regression line and few VAF_RNA_ values of 0 and 1; these patterns were seen in the majority of the trans-ReQTL correlations.

### 3.3. Overlap with eQTLs

First, we assessed the overlap between cis-ReQTLs and significant cis-eQTLs reported in the GTEx database V7 (V7 trans-eQTLs were not available at the time of the submission). Of the significant cis-ReQTL loci, 17,960 (84%) were reported as cis-eQTLs loci in the GTEx database V7 (Supplementary Table 8).

We then analyzed the 3411 SNV loci identified exclusively through ReQTL analysis, and compared them with respect to function, position and annotation, to the set of loci called by both ReQTL and eQTL analyses. Several differences between the two sets were obvious. First, the ReQTL-exclusive set of loci contained higher percentage of positions known to be subject to RNA editing (3.4% vs 0.2% in the eQTLs, p<0.0001, chi-square test). Examples of cis-ReQTLs in RNA-editing sites are shown in Figure 4a. Overall, these correlation plots were very similar, with few VAF_RNA_ values of 0 or 1 (see Figure 3c). The remaining ReQTL-exclusive SNVs showed various types of correlation patterns including some similar to eQTLs, with no obvious associations between a given type of correlation and a functional/positional annotation (Figure 4b). Second, the ReQTL-exclusive positions more often resided in non-coding exons and intergenic regions, while those identifiable by eQTLs were more frequently found in the 3’-UTR of the corresponding gene (Figure 4c, p<0.0001, chi-square test). The remaining functional categories did not substantially differ between the ReQTL-exclusive loci and those identified by both ReQTL and eQTL analysis. Additional factors, such as genome reference version (hg38, 2% of the ReQTL exclusive positions were not convertible to hg19), SNV-aware alignment, and different processing tools/versions, as well as covariate adjustments, are likely to partially account for the differences in the sets of variants identified by ReQTL and those in the GTEx database.

**Figure 4.**
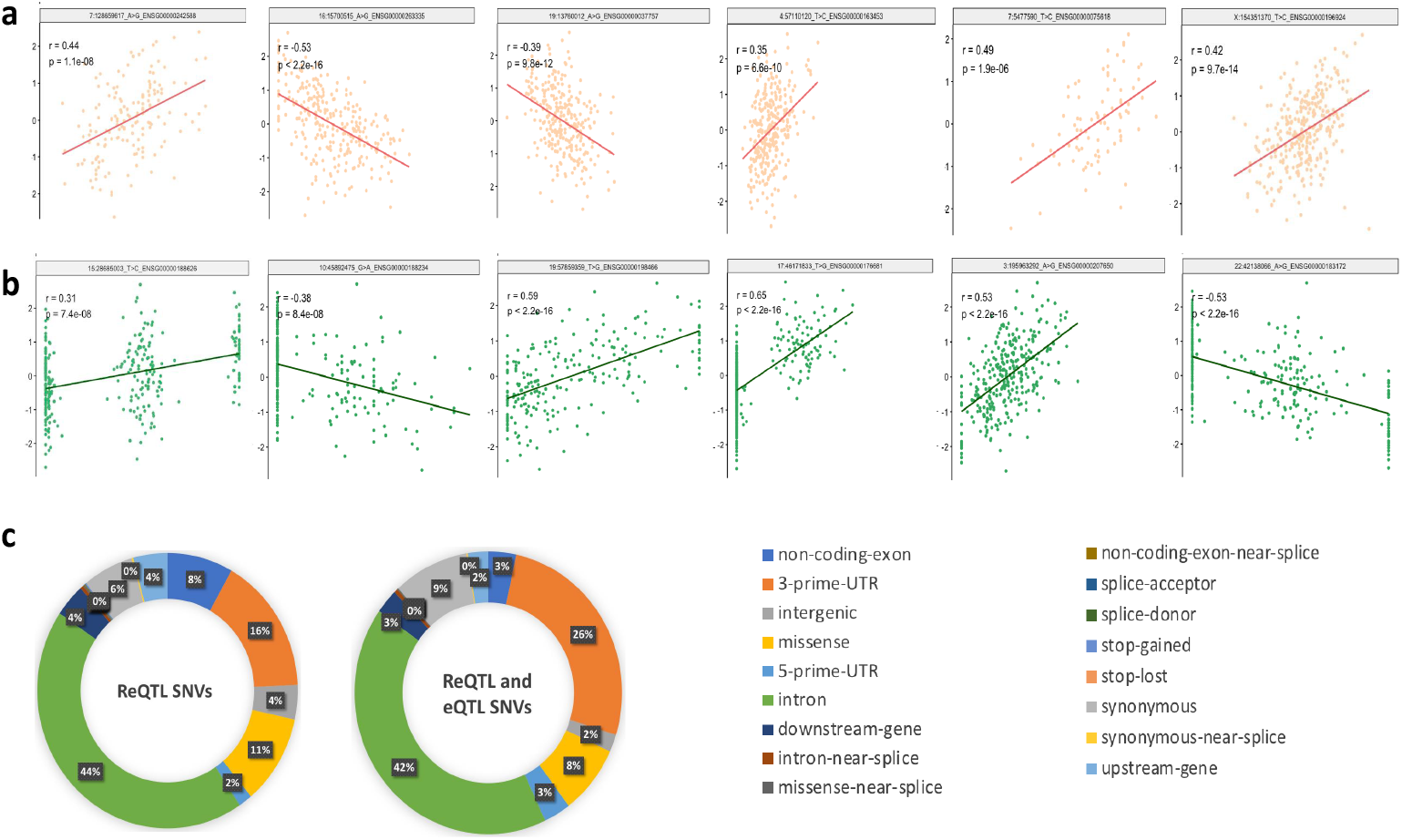
**a.** Correlations identified exclusively through ReQTL (and not through eQTL) analysis, and involving loci residing in known RNA-editing positions; all ReQTLs show similar correlation plot patterns, with few extreme VAF_RNA_ values (0 or 1). **b.** ReQTL-exclusive SNVs residing outside known RNA-editing sites; non-coding exon (left two plots), missense (middle two plots) and intergenic (right two plots) SNVs are shown; various correlation plots are seen, including eQTL-resembling ones. No association between type of correlation plot and functional/positional annotation of the SNV is observed. **c.** Distribution of the func-tional/positional annotation of the ReQTL-exclusive SNVs, as compared to eQTL-identifiable SNVs; the largest differences are seen in the distribution of non-coding, intergenic (more frequent) and missense (less frequent) SNVs involved in ReQTL-exclusive correlations.

### 3.4. Overlap with RNA-editing sites

Next, we set to assess the ReQTL capacity to capture correlations between gene expression and genetic variation introduced through the process of RNA-editing. To do that, we downloaded the list of genomic positions known to be subject to RNA editing from the REDIportal (srv00.recas.ba.infn.it/atlas, all tissues, (Picardi *et al.*, 2017)), and intersected it with the significant cis- and trans-ReQTLs identified through our analysis. From the 21,307 cis-ReQTLs loci, 152 (0.7%) coincided with RNA-editing sites. Strikingly, this percentage was substantially higher – 35.8% (816 out of the 2280, p<1e-16, chi-square test) – among the trans-ReQTL positions. These positions were implicated in an even higher percentage – 67.6% - of the individual trans-ReQTL correlations (Supplementary Table 9, examples of trans-ReQTLs in and outside RNA-editing sites are shown on Figure 5). Thus, the majority of the significant trans-correlations in our dataset involved an RNA-editing site.

**Figure 5.**
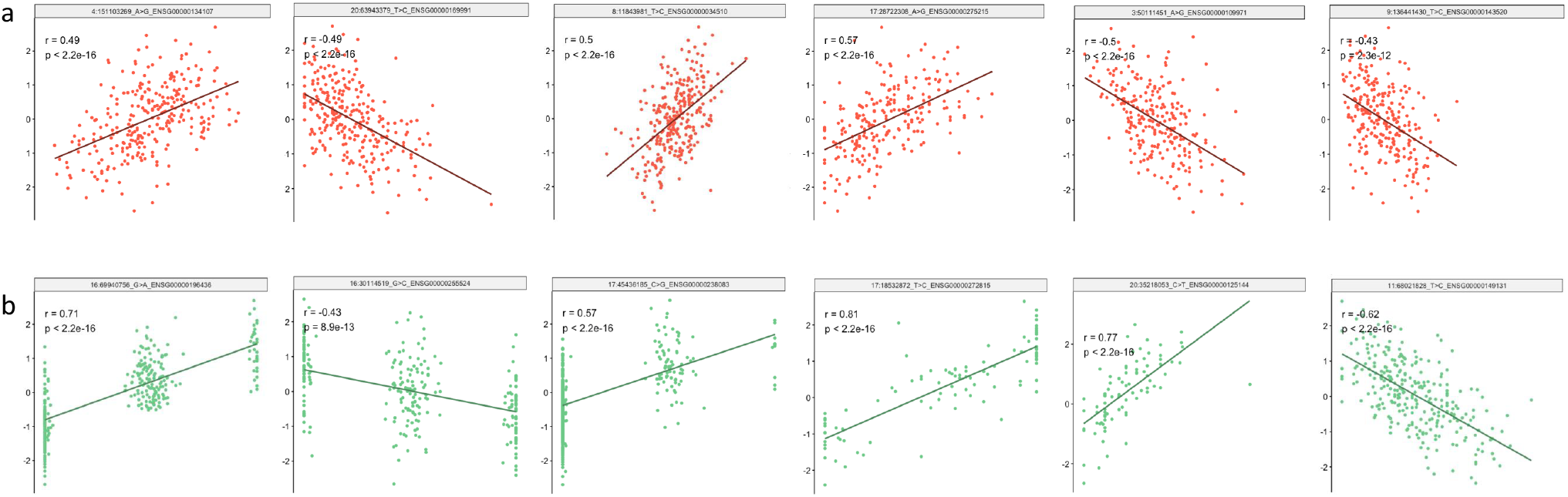
**a.** Trans-ReQTL correlations involving RNA-editing sites; all correlation plots had similar characteristic patterns. **B.** Trans-ReQTLs outside RNA-editing sites – different patterns of correlation plots were observed.

### 3.5. ReQTL usage

We note several considerations for the usage of the ReQTLs. First, because ReQTLs are based on VAF_RNA_, they are confined to expressed SNV loci in the studied sample-set and would not capture variants in transcriptionally silent genomic regions. Related to that, SNV loci with low expression levels (below the required threshold for minimum number of RNA-sequencing reads) are not eligible for this type of analysis. The threshold for minimum RNA-sequencing reads is necessary for distinguishing SNV positions with monoallelic reference signal from non-expressed SNV positions and is critical for the estimation of VAF_RNA_. In the presented results, we have selected a threshold of 10 RNA-sequencing reads covering each SNV position of interest, based on considerations for sequencing depth and confidence of the VAF_RNA_ assessment, and also following trends in the assessment of RNA editing from RNA-sequencing data (Park *et al.*, 2017). Our experiments with various minimum thresholds show that higher thresholds increase the accuracy of the VAF_RNA_ estimation, naturally retaining a lower number of variants for analysis (Movassagh *et al.*, 2016). In the readCounts package, this threshold is flexible and can be set at the desired level depending on the depth of sequencing and required confidence in the assessment of the VAF_RNA_.

With respect to gene expression, ReQTL analysis can employ data processing identical to that of eQTL, including adjustment for covariates. In this study, we closely followed the pipeline employed by the GTEx Consortium, correcting for reported age, race, sex and hidden confounders using the top 25 PEER factors based on sample size (200-300 samples) (Aguet *et al.*, 2017a). In addition, we quantile-transformed the gene expression, as is customary in eQTL analyses. As a result, we observed a strong linear correlation between quantile-transformed, covariate-adjusted gene expression and VAR_RNA._ To fully explore ReQTLs, other expression-transformation strategies (Palowitch *et al.*, 2018) may also be applicable.

VAF_RNA_ estimation can be also affected by technical parameters or settings. To minimize such effects, we apply highly conservative settings to the alignment, variant call and the read count assessment, correct for a high number of hidden confounders (top 10 VAF_RNA_ PCs), and closely follow the best practices for data processing in allelic analysis (Castel *et al.*, 2015). We observe strong concordance between the VAF_RNA_ of multiple SNVs in the same gene (Figure 6), which indicates consistency in VAF_RNA_ estimation. In addition to the growing reliability of the identification of genetic variants from RNA-sequencing data (Piskol *et al.*, 2013; Deelen *et al.*, 2015), it is important to note that ReQTLs do not necessarily require prior variant calls and can be run on custom pre-defined lists of genomic positions such as those in dbSNP or the database of RNA-editing sites. An additional factor for the VAF_RNA_ estimation is the allele mapping bias. While shown to have little to no effect on the gene expression estimation (Panousis *et al.*, 2014), mapping bias can lead to overestimation of the reference allele frequency, and consequently bias in the VAF_RNA_ assessment (Brandt *et al.*, 2015). To correct for that, we map the raw sequencing reads against a genome index containing the SNVs subjected to our analysis. As a result, in the SNV-aware mapped sequencing data, we do not observe signs of mapping bias (See Figures 3-5). Alternative strategies for mapping bias correction, including read count assessment prior to mapping, are also possible (Miao *et al.*, 2018).

**Figure 6.**
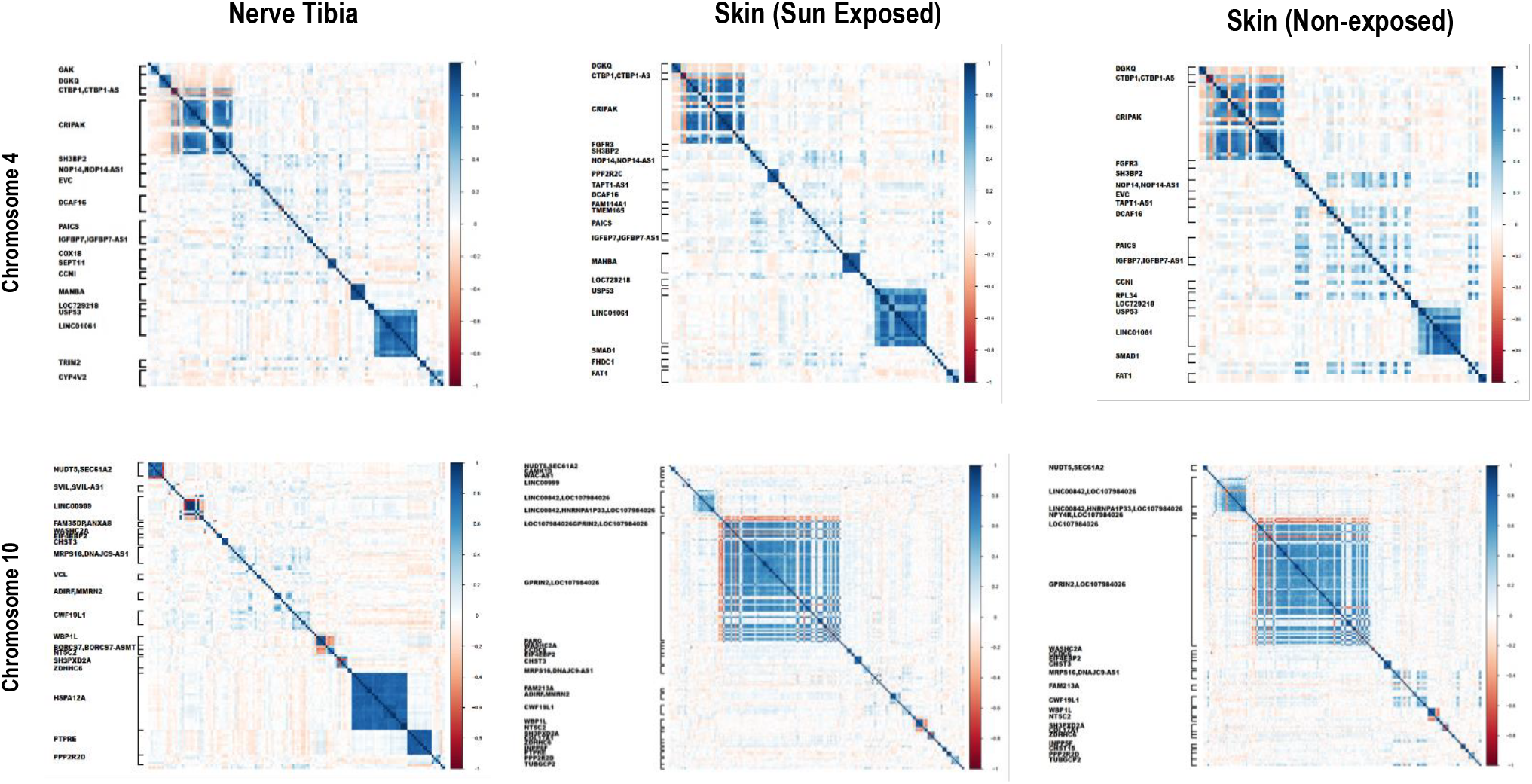
VAF_RNA_ – VAF_RNA_ correlations along genes with multiple significant SNVs within the same gene (Chromosome 4, top, Chromosome 10, bottom) in nerve (left), skin-sun-exposed (middle), and skin-non-exposed (right). Strong positive (blue, in-phase), or negative (red, inverted phase) correlations are seen between the SNV positions residing in the same gene, indicating consistency in the VAF_RNA_ estimation.

## 4. Discussion

Traditional eQTLs assess the number of variant-harboring alleles (N ∈ {0,1,2} for diploid genomes), in correlation with gene or transcript abundance across a population of individuals/samples. In our method – ReQTLs – the DNA allele count is substituted for the VAF_RNA_ at expressed SNV loci in the transcriptome; both VAF_RNA_ and the gene expression are assessed from the same sets of RNA-sequencing data.

At expressed loci, the difference between DNA- and RNA-allele counts is driven by two major mechanisms - allele-preferential expression and RNA-editing. By using RNA-allele counts, ReQTLs are readily applicable to study both of the above mechanisms and to help elucidate their increasingly recognized role in diverse cellular processes (Imprialou *et al.*, 2017; Casamassimi *et al.*, 2017; Do *et al.*, 2017; Eisenberg and Erez Y Levanon, 2018; Gagnidze *et al.*, 2018; Moreno-Moral *et al.*, 2017; Vandiedonck, 2018). Furthermore, compared to DNA-allele count, correlation of variant RNA-allele frequency with gene expression holds several technical advantages. First, as mentioned above, VAF_RNA_ constitutes a continuous measure and allows for precise quantitation of the allele representation. Second, since VAF_RNA_ and gene expression levels can be retrieved from a single source of transcriptome sequencing data alone, ReQTL analyses naturally employ within-sample concordance of the measurements. Related to the above, ReQTLs do not require matched DNA data, and are thus directly applicable to the growing amount of transcriptome sequencing data across species and conditions.

Along with the eQTL analysis on GTEx datasets, our ReQTL analysis identified a substantially higher number of cis- (as compared to trans-) correlations, the majority of which overlap with previously reported eQTL loci (Aguet *et al.*, 2017b). A notable finding on the ReQTL-exclusive loci is the high frequency of RNA-editing sites, correlated with gene expression in both cis- and trans-fashion. In our data, RNA-editing sites accounted for more than a third of the trans ReQTL loci and were involved in approximately two thirds of the trans-ReQTLs correlations, which presented with a characteristic correlation pattern (see Figures 3-5). These observations indicate the need for systematic evaluation of RNA-editing mediated effects on gene expression, for which ReQTLs can provide unique insights. In addition to RNA editing, correlations identified exclusively through ReQTL analysis (and not through eQTL analysis) are likely to include solely RNA-mediated (as opposed to DNA-mediated) molecular interactions.

In conclusion, our results show that ReQTL analysis: (1) is computationally feasible and can identify variation-expression relationships from RNA-sequencing data, (2) identifies a substantial subset of the eQTL-identifiable variants, and (3) identifies additional SNV loci which, in the herein presented results, are enriched in RNA-editing sites. Given the quickly growing availability of RNA-sequencing data, ReQTL analyses have considerable potential to facilitate the discovery of novel molecular interactions.

## References

Aguet,F. et al. (2017b) Genetic effects on gene expression across human tissues. Nature, 550, 204–213.

Albert,F.W. and Kruglyak,L. (2015) The role of regulatory variation in complex traits and disease. Nat. Rev. Genet., 16, 197–212.

De Almeida,C. et al. (2018) RNA uridylation: a key posttranscriptional modification shaping the coding and noncoding transcriptome. Wiley Interdiscip. Rev. RNA, 9.

Atak,Z.K. et al. (2013) Comprehensive Analysis of Transcriptome Variation Uncovers Known and Novel Driver Events in T-Cell Acute Lymphoblastic Leukemia. 9.

Van der Auwera,G.A. et al. (2013) From FastQ data to high confidence variant calls: the Genome Analysis Toolkit best practices pipeline. Curr. Protoc. Bioinforma., 43, 11.10.1–33.

Brandt,D.Y.C. et al. (2015) Mapping Bias Overestimates Reference Allele Frequencies at the HLA Genes in the 1000 Genomes Project Phase I Data. G3 (Bethesda)., 5, 931–41.

Brandt,M. and Lappalainen,T. (2017) SnapShot: Discovering Genetic Regulatory Variants by QTL Analysis. Cell.

Casamassimi,A. et al. (2017) Transcriptome Profiling in Human Diseases: New Advances and Perspectives. Int. J. Mol. Sci., 18.

Castel,S.E. et al. (2015) Tools and best practices for data processing in allelic expression analysis. Genome Biol., 16.

Chess,A. (2016) Monoallelic Gene Expression in Mammals. Annu. Rev. Genet., 50, 317–327.

Deelen,P. et al. (2015) Calling genotypes from public RNA-sequencing data enables identification of genetic variants that affect gene-expression levels. Genome Med., 7, 30.

Do,C. et al. (2017) Genetic-epigenetic interactions in cis: a major focus in the post-GWAS era. Genome Biol., 18, 120.

Eisenberg,E. and Levanon,E.Y. (2018) A-to-I RNA editing - immune protector and transcriptome diversifier. Nat. Rev. Genet., 19, 473–490.

Eisenberg,E. and Levanon,E.Y. (2018) A-to-I RNA editing - Immune protector and transcriptome diversifier. Nat. Rev. Genet., 19, 473–490.

Gagnidze,K. et al. (2018) A New Chapter in Genetic Medicine: RNA Editing and its Role in Disease Pathogenesis. Trends Mol. Med., 24, 294–303.

Guo,Y. et al. (2018) Single-nucleotide variants in human RNA: RNA editing and beyond. Brief. Funct. Genomics.

Heinig,M. (2018) Using Gene Expression to Annotate Cardiovascular GWAS Loci. Front. Cardiovasc. Med., 5, 59.

Imprialou,M. et al. (2017) Expression QTLs Mapping and Analysis: A Bayesian Perspective. Methods Mol. Biol., 1488, 189–215.

Kim,D. et al. (2015) HISAT: a fast spliced aligner with low memory requirements. Nat. Methods, 12.

Ko,Y.-A. et al. (2017) Genetic-Variation-Driven Gene-Expression Changes Highlight Genes with Important Functions for Kidney Disease. Am. J. Hum. Genet., 100, 940–953.

Li,H. et al. (2015) eQTL networks unveil enriched mRNA master integrators downstream of complex disease-associated SNPs. J. Biomed. Inform., 58, 226–234.

Miao,Z. et al. (2018) ASElux: an ultra-fast and accurate allelic reads counter. Bioinformatics, 34, 1313–1320.

Moreno-Moral,A. et al. (2017) Systems Genetics as a Tool to Identify Master Genetic Regulators in Complex Disease. Methods Mol. Biol., 1488, 337–362.

Movassagh,M. et al. (2016) RNA2DNAlign: nucleotide resolution allele asymmetries through quantitative assessment of RNA and DNA paired sequencing data. Nucleic Acids Res., 44.

Odhams,C.A. et al. (2017) Mapping eQTLs with RNA-seq reveals novel susceptibility genes, non-coding RNAs and alternative-splicing events in systemic lupus erythematosus. Hum. Mol. Genet., 26, 1003–1017.

Palowitch,J. et al. (2018) Estimation of cis-eQTL effect sizes using a log of linear model. Biometrics, 74, 616–625.

Panousis,N.I. et al. (2014) Allelic mapping bias in RNA-sequencing is not a major confounder in eQTL studies. Genome Biol., 15, 467.

Park,E. et al. (2017) Population and allelic variation of A-to-I RNA editing in human transcriptomes. Genome Biol.

Picardi,E. et al. (2017) REDIportal: A comprehensive database of A-to-I RNA editing events in humans. Nucleic Acids Res.

Piskol,R. et al. (2013) Reliable identification of genomic variants from RNA-seq data. Am. J. Hum. Genet., 93, 641–51.

Shabalin,A.A. (2012) Matrix eQTL: Ultra fast eQTL analysis via large matrix operations. Bioinformatics, 28, 1353–1358.

Stegle,O. et al. (2012) Using probabilistic estimation of expression residuals (PEER) to obtain increased power and interpretability of gene expression analyses. Nat. Protoc., 7, 500–507.

Vandiedonck,C. (2018) Genetic association of molecular traits: A help to identify causative variants in complex diseases. Clin. Genet., 93, 520–532.

Weiser,M. et al. (2014) Novel distal eQTL analysis demonstrates effect of population genetic architecture on detecting and interpreting associations. Genetics.

Winter,J.M. et al. (2018) Modifier locus mapping of a transgenic F2 mouse population identifies CCDC115 as a novel aggressive prostate cancer modifier gene in humans. BMC Genomics, 19, 450.

